# *qFit-ligand* reveals widespread conformational heterogeneity of drug-like molecules in X-ray electron density maps

**DOI:** 10.1101/253419

**Authors:** Gydo C.P. van Zundert, Brandi M. Hudson, Daniel A. Keedy, Rasmus Fonseca, Amelie Heliou, Pooja Suresh, Kenneth Borrelli, Tyler Day, James S. Fraser, Henry van den Bedem

## Abstract

Proteins and ligands sample a conformational ensemble that governs molecular recognition, activity, and dissociation. In structure-based drug design, access to this conformational ensemble is critical to understand the balance between entropy and enthalpy in lead optimization. However, ligand conformational heterogeneity is currently severely underreported in crystal structures in the Protein Data Bank, owing in part to a lack of automated and unbiased procedures to model an ensemble of protein-ligand states into X-ray data. Here, we designed a computational method, *qFit-ligand*, to automatically resolve conformationally averaged ligand heterogeneity in crystal structures, and applied it to a large set of protein receptor-ligand complexes. We found that up to 29 % of a dataset of protein crystal structures bound with drug-like molecules present evidence of unmodeled, averaged, relatively isoenergetic conformations in ligand-receptor interactions. In many retrospective cases, these alternate conformations were adventitiously exploited to guide compound design, resulting in improved potency or selectivity. Combining *qFit-ligand* with high-throughput screening or multi-temperature crystallography could therefore augment the structure-based drug design toolbox.

## Introduction

Ligands and their protein receptors sample an ensemble of conformations in solution. The energetic contribution of conformational entropy plays a critical role in receptor-ligand molecular recognition,^1,2^ but precisely how distinct conformations contribute to binding, activity, and dissociation often remains poorly characterized. The majority of 3-dimensional protein structures are single, static models obtained from X-ray crystallography by averaging over the unique conformations in the unit cells. Crystallographic B-factors quantify harmonic displacements from average atomic positions, but are adversely affected when unmodeled discrete alternate conformations overlap. By their nature, such static, harmonic models cannot rationalize molecular attributes that rely on dynamic, anharmonic displacements of atoms.^3,4^ Revealing discrete conformations^5^ that are more fully representative of the receptor-ligand conformational ensemble from X-ray electron density maps would overcome this limitation, and create new opportunities to address open questions in chemical and structural biology. For example, such models might help to provide a structural basis for on-pathway intermediates in substrate binding or release detected by NMR.^6,7^

Additionally, an incomplete picture of receptor-ligand structural dynamics impedes structure-based drug design. While overall binding affinities measured in solution report on a receptor-ligand ensemble, the structure-activity relationships are often informed by static models for further optimization. During small molecule optimization, even minor chemical changes can lead to altered binding modes unforeseen from the static models, frustrating design^8,9^ (**Figure 1A,B,C**). Examples where subtle changes in chemical structure of the ligand led to different binding modes are abundant. Fragment optimization of CDK8 inhibitors revealed that small modifications led to a new binding mode, which was exploited to develop potent and selective inhibitors.^10^ Dramatically different binding modes as a result of minor changes in the chemical structure of the ligand are also illustrated by Hsp90 and PTR1 inhibitors. In the course of structure-based optimization of Hsp90 inhibitors, Casale and coworkers observed a flipped binding mode of the ligand in the crystal structure, leading them to change the direction of design towards a low nanomolar compound (**Figure 1B**).^11^ In PTR1, compounds that presumably could not be accommodated by the binding site ultimately led to a boost in affinity, owing to altered binding modes.^11,12^ In human lipoprotein-associated phospholipase A2, two related ligands (**Figure 1C**) that explored distinct subpockets were merged into a more potent ligand (**Figure 1D**).^13^ These selected examples highlight both the perils of design based solely on an initial ligand pose, but also how the fortuitous discovery of alternate poses can create new opportunities for design.

**Figure 1.**
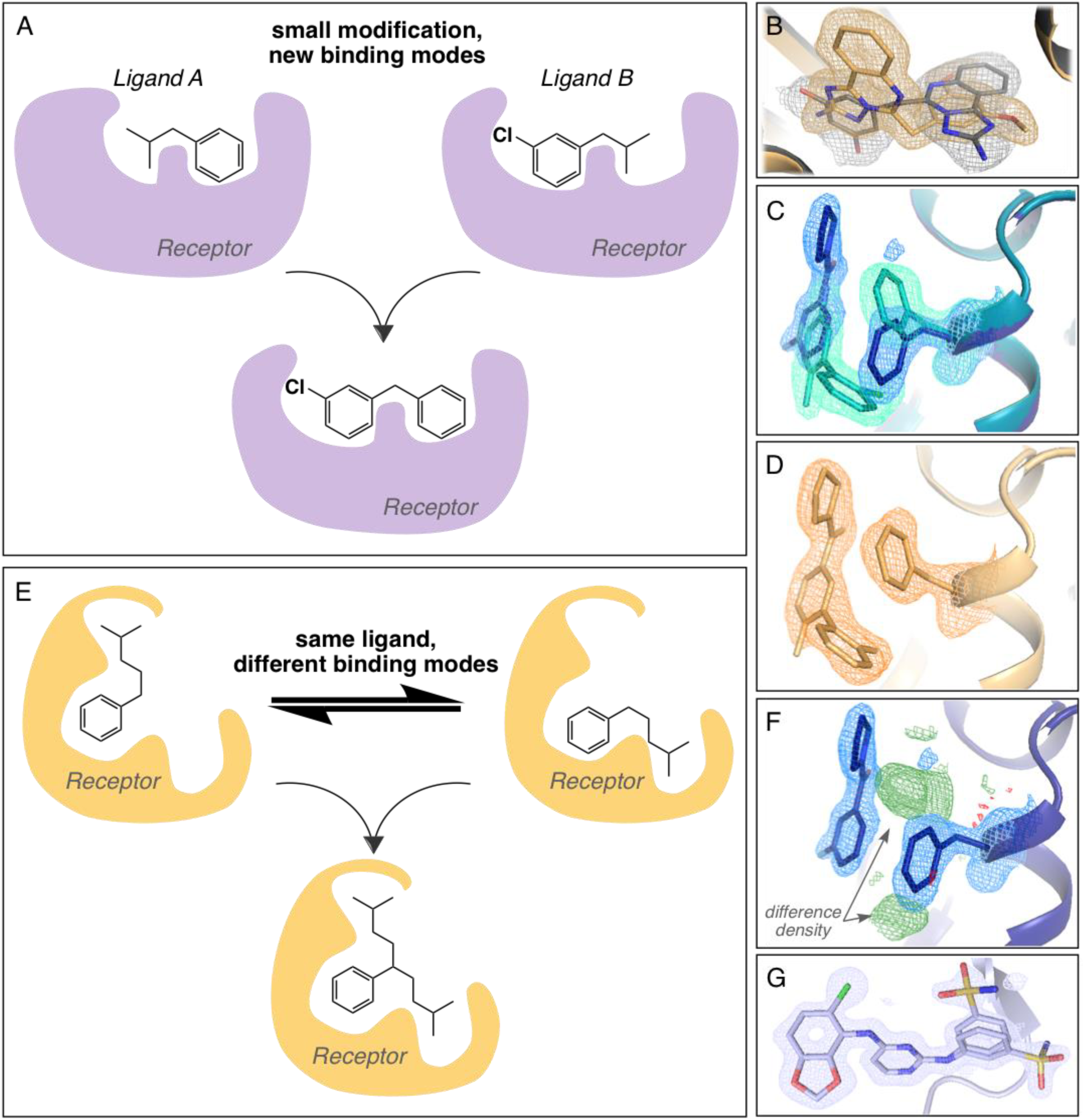
Ligand structural dynamics and minor changes during fragment optimization lead to new binding modes and drive drug design. A) Subtle changes in chemical structure of ligands can impose new binding modes. B) Hsp90 inhibitors in gray (PDB ID 4CWO) and gold (PDB ID 4CWN). C) Subtle changes in chemical structures lead to changes in binding pose for Lp-PLA2, changing the course of design (PDB IDs 5JAL, 5JAO). D) New Lp-PLA2 inhibitor designed as a result of observed alternate binding poses of fragments (PDB ID 5JAP). E) Near isoenergetic receptor-ligand conformations exchange in dynamic equilibrium in crystal structures. These conformations can inform the design of a ligand with higher affinity. F) Evidence of difference density in x-ray crystal structure of Lp-PLA2 fragment shows alternate binding poses pre-exist at low occupancy (PDB ID 5JAL). G) Alternate conformations exploited in the design of EphB4-binding ligands (PDB ID 2VWX). Electron densities are shown at 1.5σ. Positive (green) and negative (red) difference densities in panel J are shown at +3.0σ and −3.0σ, respectively.

One hypothesis to explain how subtle modifications cause a switch to a second binding pose is that the unmodified ligand samples the second pose at low, but potentially detectable, occupancy (**Figure 1E**). Indeed, in the phospholipase example above, the presence of difference electron density suggested that multiple conformations might have been sampled in at least one of the smaller ligands (**Figure 1F**). This view posits that degeneracies in ligand binding modes can also be accessed by small modifications of the ligand chemical structure that shift the receptor-ligand equilibrium ensemble. This hypothesis is supported by anecdotal examples, such as long time scale molecular dynamics (MD) simulations of trypsin, in which both the ligand and receptor adopt many stable configurations.^14^ Experimentally, multiple conformations can be present in X-ray electron density, as in HDAC6 where many related ligands were modeled in multiple conformations in concert with distinct conformations of the receptor.^15^ In other cases where the alternate conformations have been experimentally revealed, they have been exploited to improve affinity. For example, X-ray crystallography revealed alternate conformations for singly-substituted EphB4 ligands that inspired the creation of bis-substituted ligands with increased potency (**Figure 1G**).^16^ NMR measurements of conformational heterogeneity for ligands generated against the antibacterial target LpxC uncovered a larger cryptic envelope that was filled by larger, more potent ligands.^17^ Collectively these examples suggest that multiple ligand poses are likely energetically accessible for many proteins. Fully realizing the potential of this phenomenon in structure-based design, exemplified by EphB4 and LpxC inhibitors, requires reliable characterization of pre-existing conformational heterogeneity (**Figure 1E**) of ligand-receptor complexes.

While protein conformational heterogeneity has been automatically and systematically characterized in X-ray crystallography data,^18–22^ ligand conformational heterogeneity is less explored. Various software algorithms can identify and build ligands into electron density maps without human intervention;^23–28^ however, these approaches typically provide several top scoring ligand conformations at unit occupancy. While in principle the user can select multiple candidate conformations for the final model, none of these approaches consider an ensemble of alternate conformations from the outset. Moreover, they may build unrealistic conformations that incorrectly fit into the ensemble-averaged electron density.

Here, we present a new, automated approach based on *qFit*,^19,20^ called *qFit-ligand*, to create parsimonious multiconformer ligand models in crystallographic electron densities. We first surveyed the PDB to investigate ligand heterogeneity in current crystal structures, and selected a diverse, curated benchmark set of pharmaceutically relevant protein targets with alternate ligand conformations across a wide range of resolution and occupancy (**Table S1**). We found that *qFit-ligand* can detect alternate conformations at occupancies down to 20 %, even at relatively modest resolutions of 2.0Å. We then applied our method prospectively to all cases of the *Drug Design Data Resource* (*D3R*, *drugdesigndata.org*), a subset of the *Twilight database,^29^* and all PDB entries for the bromodomain-containing protein 4 (BRD4), revealing unmodeled alternate conformations in 29 % of the cases. To evaluate the quality of our multiconformer ligands, we calculated R-factors and ligand energies relative to a single conformer ligand model. Our results indicate that *qFit-ligand* is a powerful, efficient, and user-friendly tool to model and discover alternate ligand conformations.

## Results

### Creating a benchmark set of true positive ligand alternative conformations from the PDB

To estimate the prevalence of multiconformer ligands in crystal structures, we surveyed all 130,054 PDB entries as of June 2017 that contained non-covalently bound ligands with more than 15 non-hydrogen atoms in their X-ray crystal structure. This resulted in 44,620 PDB entries totaling 133,724 ligands. Of those, 2,611 ligands, or less than two percent (1,078 unique ligand codes), distributed over 1,845 PDB entries consisted of two or more alternate conformations (Materials and Methods, **Figure S1**). Many of these molecules are common crystallographic additives (PEG, cholesterol, etc.) or metabolites (ATP, NADPH, etc.). We therefore manually curated a true positive benchmark set of receptors of pharmaceutical interest containing multiconformer, drug-like molecules. Cases where ligands adopted entirely different binding modes, such as flipped ligands, were discarded. This resulted in 90 crystal structures that could be stably refined against the deposited structure factors and CIF restraints files (**Table S1**).

We apportioned the conformational heterogeneity of ligands in our benchmark set into four categories (**Figure 2A**): terminal end flips, where only terminal atoms are flipped/rotated; ring flips where a ring system is flipped, usually by 180 degrees; branching ligands, where a side-chain or branch of a ligand has an alternate conformation; and displaced ligands, where all atoms are at least slightly displaced in combination with differences in their internal degrees of freedom. The benchmark set heavily overrepresented 50/50 occupancy splits (**Figure 2B**), reflecting a historical tendency against refining occupancies in favor of refining B-factors only; however, re-refinement with Phenix^30^ substantially broadened the distribution (**Figure 2B**). Unlike the occupancies, the RMSD between alternate conformations was similar after re-refinement (**Figure 2C**). Interestingly, we observed no correlation between occupancy shift and difference in mean B-factor (**Figure 2D**), in contrast to earlier reports.^31,32^

**Figure 2.**
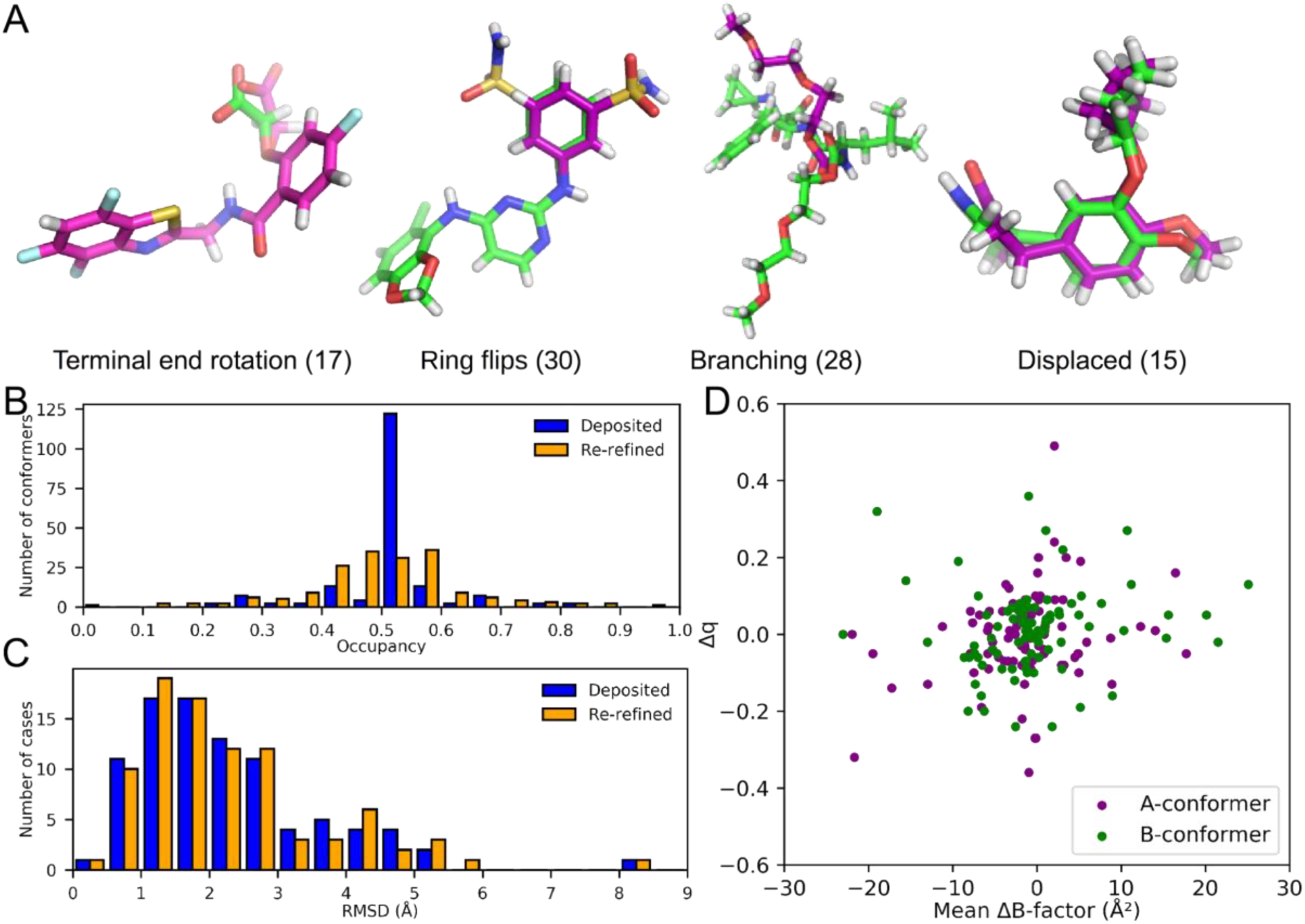
Benchmark statistics. A) Categories of alternate conformations present in the benchmark. B) Conformer occupancies pre‐ (blue) and post‐ (orange) re-refinement. C) Ligand A to B conformer RMSD, pre (blue) and post (orange) re-refinement. D) Occupancy shift versus mean B-factor difference after re-refinement.

### Developing the qFit-ligand algorithm and calibration against synthetic data

We designed the *qFit-ligand* algorithm to iteratively explore a vast conformational space to determine a parsimonious ensemble of up to five occupancy-weighted conformations that, collectively, optimally fits the electron density (**Figure 3A**, Materials and Methods). Briefly, *qFit-ligand* takes as input a refined, single conformation receptor-ligand structure in PDB format, and a 2mF_o_-DF_c_ density map. It first determines rotatable bonds and rigid groups of atoms within the ligand. Starting from each rigid group, *qFit-ligand* performs a local, six-dimensional translational and rotational search in the rigid group’s neighborhood, selecting up to five occupancy-weighted candidate positions that, collectively, minimize the real-space density residual of the rigid group. In subsequent steps, *qFit-ligand* iteratively grows the rigid group by exhaustively sampling increments of several torsion angles simultaneously, while avoiding collisions with the receptor. At each step, it selects up to five occupancy-weighted conformations by again minimizing the density residual. This is repeated until all torsion angles are determined and the full ligand is built up. The maximum number of conformations generated by *qFit-ligand* at this stage is five times the number of rigid groups. The final occupancy-weighted, multiconformer ensemble is selected from this pool by combining cross-correlation, geometric, and density residual measures. The ligand multiconformer ensemble is then combined with the receptor and refined with phenix.refine.

**Figure 3.**
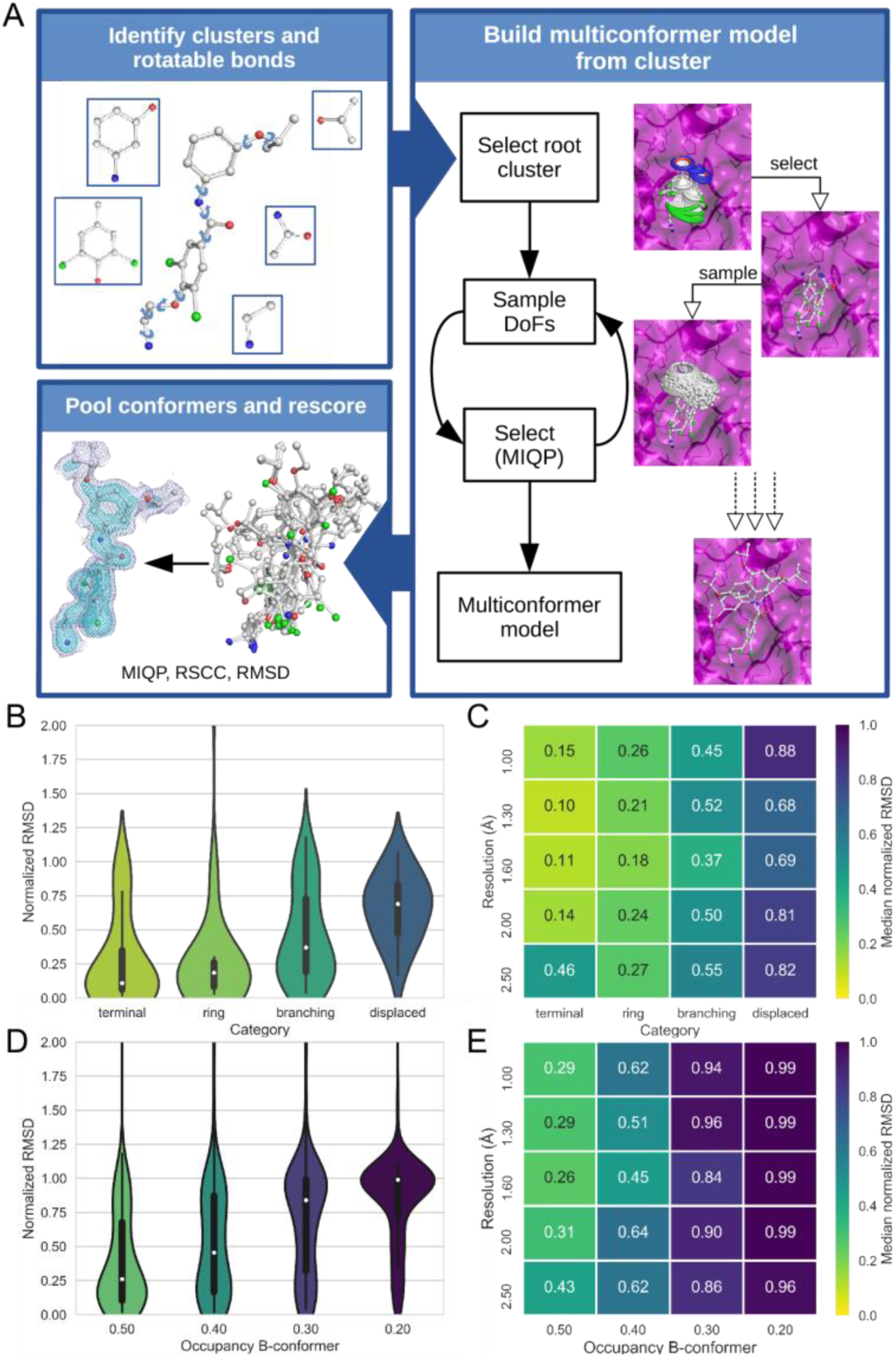
*qFit-ligand* workflow and statistics on synthetic data. A) Rigid clusters are defined as rings or terminal ends, and rotatable bonds as any bond that is not part of a ring. The local search finds possible positions and orientations of each cluster in the binding site, avoiding steric clashes with the protein. As clusters are joined, torsions and degrees of freedom are sampled. Up to 5 ligand conformations that best match the ligand density are selected and combined with the protein model to give a final ligand multiconformer model. B) Violin plot of categories at 1.60Å resolution and occupancy 0.50 across 90 test cases. The white dot represents the median, the bold center line represent the interquartile range (IQR), and the thin center line represents the percentile range 25^th^-1.5 IQR to 75^th^+1.5 IQR. C) Heatmap of category vs. resolution at 0.50 occupancy. D) Violin plot for representative resolution 1.60Å and 0.50 / 0.50 occupancy at optimal parameters. E) Heatmap of normalized RMSD at optimal parameters (resolution vs. occupancy).

We calibrated *qFit-ligand* on synthetic data calculated from the benchmark set at varying resolutions and occupancies (Materials and Methods, **Figure S2**). To create starting models we deleted the alternate ‘B’ conformation of the ligand, set the occupancy of the ‘A’ conformation to 1.0, and re-refined against the deposited structure factors to reposition the ‘A’ conformation as the only modeled conformation.

We first determined optimal sampling parameters for *qFit-ligand*, balancing accuracy of the results and computational demands, measured by the minimum normalized RMSD (nRMSD, Materials and Methods) between *qFit-ligand* generated conformations and the benchmark B conformation. nRMSD values range from 0 to 1, with values closer to 0 indicating better performance; the normalization controls for the fact that the naïve RMSD between alternate conformers is higher for some ligands than for others. To compare performance we report the median minimum nRMSD of the benchmark set for different resolutions and occupancies. Our analysis suggested that sampling two torsion angles simultaneously at 6° intervals gave the best result for all resolutions and occupancies, while limiting computational costs (**Figure S3**).

Benchmarking *qFit-ligand* showed performance differences across category, resolution, and true occupancy. We evaluated the performance of *qFit-ligand* for each category of conformational heterogeneity (**Figure 3B, 3C**). At a resolution of 1.60Å and equal occupancies for the A and B conformer, the median nRMSD is 0.11 for terminal end flips, 0.18 for ring flips, 0.37 for branched ligands and 0.69 for displaced ligands (**Figure 3B**). Unsurprisingly, *qFit-ligand* performance decreased with increasing complexity of conformational heterogeneity.

The resolution dependence is more complex, however, with superior performance of *qFit-ligand* at an intermediate resolution of 1.60Å (**Figure 3C, 3E**). We attribute this to more sharply defined density peaks at high resolutions relative to intermediate resolutions. Undersampling of conformations during ligand building can result in failure to accurately hit the density peaks at high resolution, thereby leading to suboptimal scores of electron density-based measures.

The performance of *qFit-ligand* is sensitive to the occupancy of the alternate conformation, indicated by an increasing median nRMSD for lower occupancies of the B conformation (**Figure 3D, 3E**). At occupancies of 0.2 and below, the alternate conformer was rarely detected at any resolution. While *qFit-ligand* samples conformations close to the alternate conformer, evidenced by favorable (low) nRMDS before final rescoring (**Figure S3**), selecting them at low occupancies would increase the false positive rate (data not shown).

### qFit-ligand re-identifies low energy alternative conformations in experimental data

Next, we applied our method to the experimental benchmark data set with only the ‘A’ conformation retained in the re-refined starting model (**Figure 4**). *qFit-ligand* performance with real data followed the trend we observed with simulated data: localized conformational disorder like terminal end flips and ring flips were determined with higher accuracy than branched or displaced disorder. In the case of terminal ends, only one or two atoms report on the distance between alternate conformations, but the nRMSD is dominated by small coordinate shifts distributed over the entire ligand. Despite a median nRMSD of 0.56 for terminal ends, the median nRMSD for only the ‘reporting’ atoms is less than 0.30 (**Figure 4A**). Consequently, 13 out of 17 terminal end flip *qFit-ligand* results were sufficiently accurate to recognize the benchmark alternate conformer. Similarily, *qFit-ligand* determined ring rotations to within a median nRMSD of 0.28. However, the median nRMSD for branched cases is 0.69 and for displaced cases is 0.89. If we conservatively designate a *qFit-ligand* result with more than one conformation and nRMSD > 0.6 as a false positive, the false positive rates for each category are 24 % (terminal flip), 24 % (ring flip), 59 % (branching), and 87 % (displaced) (*see Discussion section regarding false positives*).

**Figure 4.**
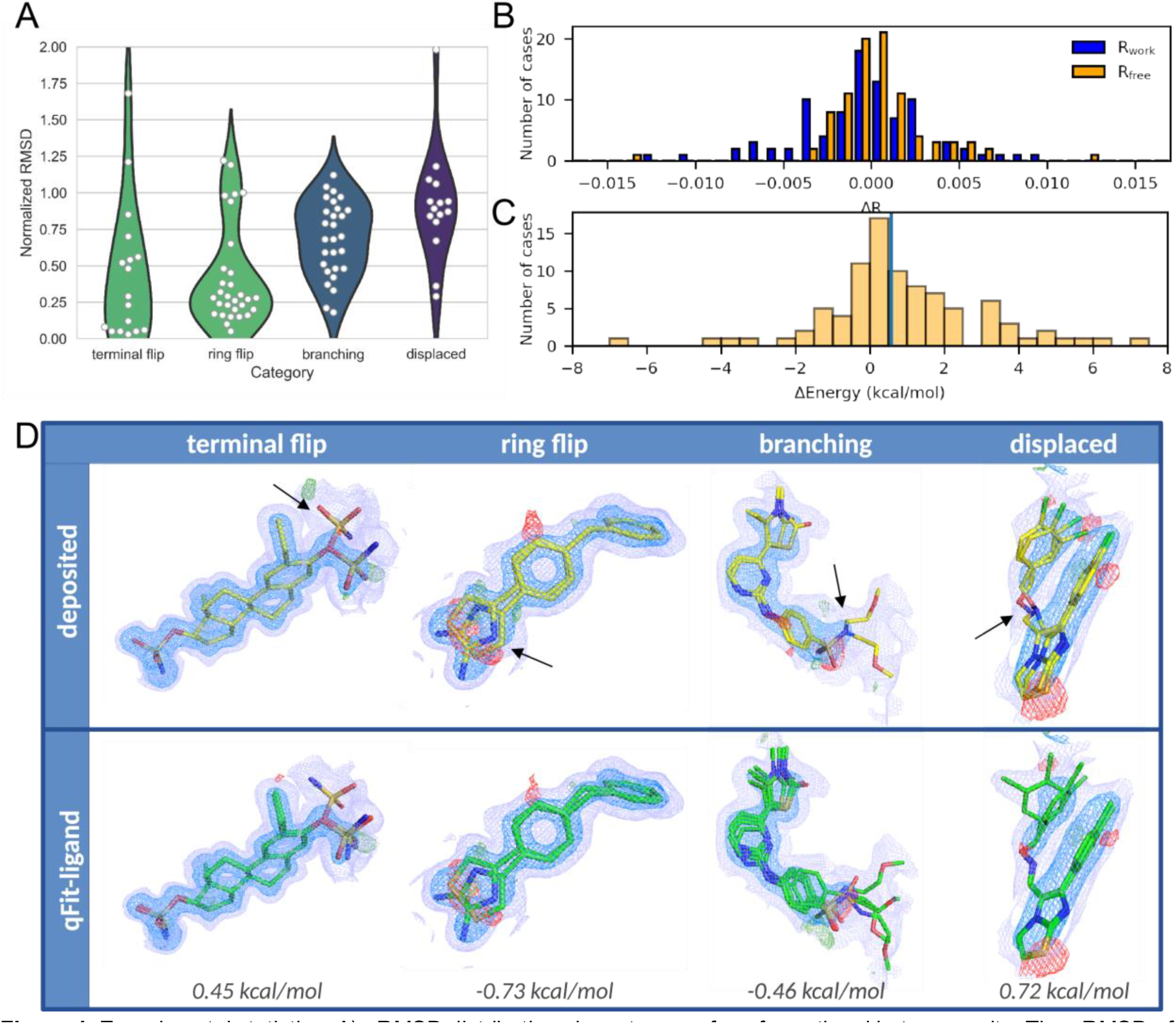
Experimental statistics. A) nRMSD distributions by category of conformational heterogeneity. The nRMSD of terminal end flips was determined from only atoms affected by the dihedral changes. B) Histogram of R_work_ and R_free_ differences between refined *qFit-ligand* models and single conformer structures. Negative values indicate a lower R for refined *qFit-ligand* models. C) The distribution of occupancy-weighted, ligand energies of *qFit-ligand* multiconformer ligands relative to single ‘A’ conformation. Negative values indicate that the multiconformer ensemble has lower internal energy than the single benchmark ‘A’ conformation. Positive values indicate the multiconformer ensemble has higher internal ligand energies. The blue horizontal line represents the mean. D) Examples of conformational heterogeneity: terminal flip (PDB ID 3OIK), ring flip (PDB ID 4L2L), branching (PDB ID 2XMY), and displaced (PDB ID 1XVP).

The distribution of R_free_ values of refined *qFit-ligand* models were nearly identical to that of the single conformer ligand models (*qFit-ligand* x̄ =0.2130, single conformer x̄ =0.2131; p-value=0.87, two-sided t-test), and statistically indistinguishable from the deposited, manually curated multiconformers (x̄ =0.2123; p-value=0.3) (**Figure 4B, Figure S4**, **Table S2**). Refined *qFit-ligand* multiconformer models improved R_free_ in 52 % of cases compared to single conformer ligand models. Although further manual refinement cycles of all 90 examples would be beyond the scope of this study, we note that elevated R_free_ values from fully automated modeling generally improve with manual refinement.

To evaluate the quality of multiconformer ligand models, we examined internal ligand energies. We found a median conformationally-averaged excess energy of 0.59 kcal/mol, i.e. the energy of ligand multiconformer models were, on average, nearly indistinguishable from that of the rerefined single ‘A’ conformation in our benchmark set (**Figure 4C**, **Figure 4D**, **Table S2**, Supplementary Material and Methods). Interestingly, automatically building a *qFit-ligand* multiconformer model in some cases substantially reduced the ligand energy compared to the single ‘A’ conformation. For example, for acyliminobenzimidazole inhibitor 36 in complex with human anaplastic lymphoma kinase,^33^ a series of concerted dihedral angle changes resulted in conformations that better fit the density, and reduced the energy by nearly 7 kcal/mol (**Figure S5**). This suggests that a ligand can accumulate strain energy when it is forced into an averaged conformation to fit the density. On the other hand, the distribution of ligand excess energies suggests that ligands also access higher energy conformations, within a few kcal/mol from the single conformation (**Figure 4D**). Indeed, ligands generally may not bind in the lowest energy conformation, or even adopt a local minimum.^34^ Favorable non-covalent interactions with the receptor, buried hydrophobic surfaces, or desolvation of ordered waters in the binding pocket can overcome penalties of strained conformations.^35,36^ Thus, alternate conformations, even at elevated ligand energies, may reduce the free energy of the receptor-ligand complex.

### qFit-ligand discovers new alternate conformations in the D3R and the Twilight Databases

For prospective discovery, we first applied *qFit-ligand* to the 145 crystal structures in the D3R dataset, a high-quality collection of manually curated protein-ligand crystal structures ranging in resolution from 1.26 to 2.75Å, designed for validation and improvement of methods in computer-aided drug design. Of the ten crystal structures in the D3R dataset with alternate ligand conformations, *qFit-ligand* recovered seven to within a median nRMSD of 0.24. Four of these overlap with our benchmark (PDB ID 4FV3, **Figure 5A**; and PDB IDs 4EK6, 4EK8, 4Y6D). We ranked all *qFit-ligand* multiconformer ligands using the Fisher z-transformation, a cross-correlation based metric which measures if alternate conformations are supported by the electron density (Materials and Methods). Three of the top four ranked ligands already had a modeled alternate conformation, and six out of seven recovered multiconformer ligands ranked within the top 20, indicating that the Fisher-z transformation is an effective ranking measure. A new alternate ligand conformation was uncovered by *qFit-ligand* in the crystal structure of the E166A mutant of *Serratia fonticola* carbapenemase (PDB ID 4EV4) from the D3R dataset (**Figure 5B**). The terminal propanol functional group of a bound meropenem intermediate adopts a previously undetected conformation. Thus, *qFit-ligand* recovered 70 % of ligand alternate conformations and even revealed a new alternate conformation in highly scrutinized experimental data.

**Figure 5.**
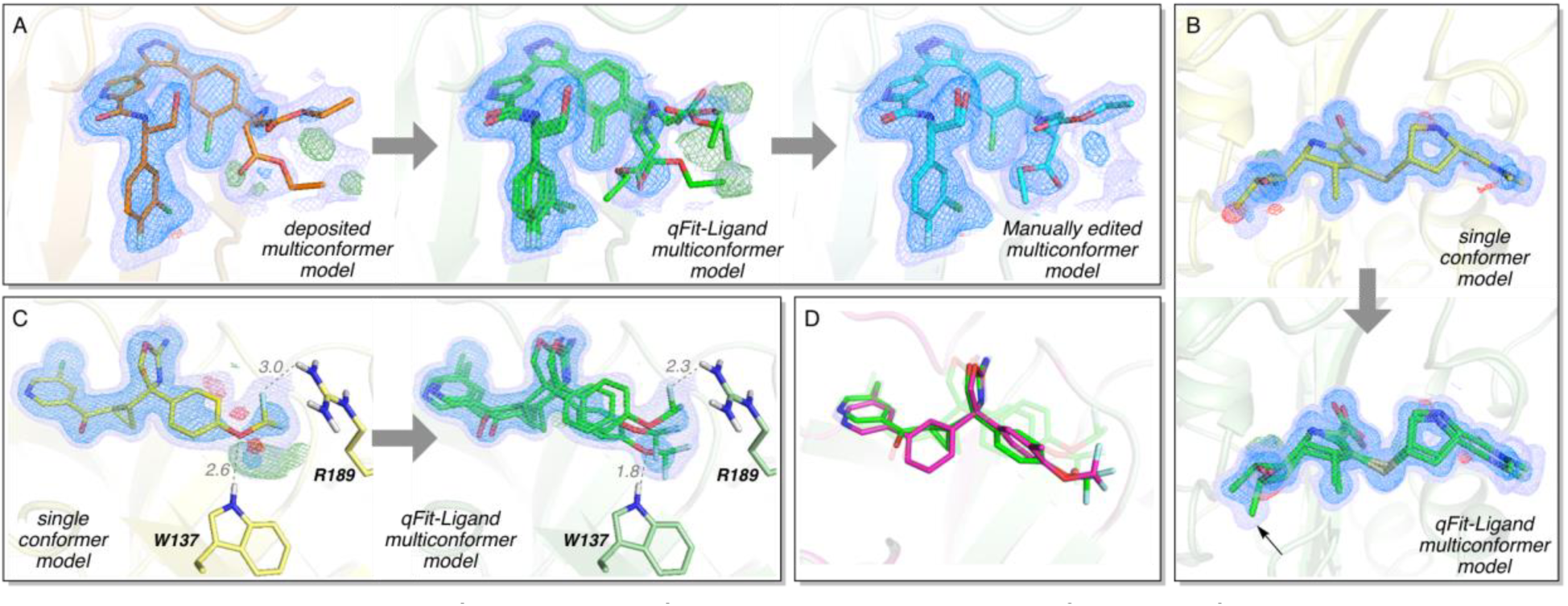
Prospective discovery of additional conformations and recovered conformations from the D3R and Twilight dataset. A) The deposited multiconformer model, *qFit-ligand* multiconformer model, and manually edited multiconformer model of ERK2 (PDB ID 4FV3). B) Single conformer and *qFit-ligand* multiconformer models of serratia fonticola carbapenemase E166A mutant with the acylenzyme intermediate of meropenem (PDB ID 4EV4). C) Prospective application of *qFit-ligand* to inhibitor 5T5 of BACE-1 (PDB ID 5EZX). Single conformer and *qFit-ligand* multiconformer models shown. D) Overlay of multiconformer model of inhibitor 5T5 (green) with inhibitor 5T6 (magenta) (PDB ID 5EZZ). Electron densities are shown at 1.5σ (blue) and 0.3σ (purple). Positive (green) and negative (red) difference densities are shown at +3.0σ and −3.0σ, respectively. All structures shown have been refined using Phenix.

We also applied *qFit-ligand* to a subset of the Twilight data set. The Twilight data set represents ligand structures in the PDB poorly supported by the electron density map, potentially indicating conformational disorder in the map, or incorrectly modeled ligands. We applied *qFit-ligand* to ligands in the Twilight database with 15 to 36 non-hydrogen atoms, resolutions better than 2.0Å, and a correlation coefficient higher than 0.6, resulting in 2,379 cases over 1,168 PDB entries, which we ranked by Fisher z-score to identify ‘hits’ of unmodeled, alternate conformations (**Table S3**). We proceeded by manually inspecting the top 10 % of cases.

In many cases, the electron density near the ligand was severely disordered, consistent with the intent of the Twilight database to flag questionable models of ligands. While *qFit-ligand* suggested alternate conformations, their validity could not unambiguously be confirmed using electron density measures. Nonetheless, in some instances significantly improved receptor-ligand interactions signified plausible alternate ligand conformations (**Figure S9, S10**).

However, for several crystal structures *qFit-ligand* unambiguously detected alternate ligand binding conformations. For example, in the crystal structure of BACE-1, *qFit-ligand* finds three conformations (occupancies of 0.32/0.36/0.32) of inhibitor 5T5 (PDB ID 5EZX) (**Figure 5C**). These conformations show the potential to engineer a ligand that can accommodate strong hydrogen bonding interactions to R189 and W137. In the single conformer model, one of the difluoromethoxy fluorines of the ligand and the amino group of R189 interact weakly through a 3.0Å hydrogen bond.^37^ Our multiconformer model shows that the ligand can adopt a position where strong hydrogen bonding between these groups occur (i.e., shorter contact distances), although at the expense of sacrificing a favorable hydrogen bonding interaction to W137. In addition, one of the *qFit-ligand* conformations turns a non-ideal hydrogen bond geometry in the crystal structure between the difluoromethoxy oxygen and amino group of W137 into an ideal geometry. Strikingly, the difluoromethoxy group of BACE-1 inhibitor 5T6 (PDB ID 5EZZ) has a binding conformation that exploits this same hydrogen bonding interaction with W137 sampled by 5T5 at low occupancy (**Figure 5D**).^38^ These insights could be used to create a new ligand with branching substituents that can simultaneously form strong hydrogen bonding interactions with R189 and W137 in hopes to increase binding affinity.

### qFit-ligand identifies widespread conformational heterogeneity in BRD4 ligand

In another notable example from the Twilight data set, *qFit-ligand* identified a minor, unmodeled population of inhibitor compound BDOIA383 bound to bromodomain-containing protein 4 (BRD4), a BET (bromodomain and extra terminal domain) BRD (**Figure 6A**, PDB ID 5CFW). BRDs are small, epigenetic ‘readers’, which recognize and bind histone acetylated lysine (AcK).^39^ BRDs can thereby epigenetically control gene transcription, and have recently emerged as important drug targets.^40^ The human genome contains 61 BRDs, distributed over 46 diverse proteins.^39^ Several potent small-molecule inhibitors for BRD4 have been structurally characterized, but designing modulators selective between BRD4 and CREB binding protein (CBP) has proved challenging.^41–43^

The BRD4-BDOIA383 example highlights how alternate conformations can expose molecular surfaces away from the primary recognition site that could be exploited for selectivity. Inspection of the major population of BDOIA383 bound to BRD4 revealed that it is stabilized by a crystal contact between the BDOIA383 morpholine oxygen and the K91 carbonyl (**Figure 6A**). By contrast, the minor conformation does not engage in crystal contacts, and buries nearly 6 % more ligand solvent accessible surface area (3115 Å^2^) than the major conformation (3302 Å^2^) (**Figure 6B**). Additionally, the minor conformation interacts with D145 at the N-terminus of helix *α*C (**Figure 6A,B**). Interestingly, compound BDOIA383 in complex with CBP BRD revealed a rotation of the isoxazole-benzimidazole bond by 180°, exposing the phenethyl group to substituted R1173 at the *α*C N-terminus^42^ (PDB ID 5CGP), occupying the space of the *qFit-ligand* minor morpholine conformation in BRD4. Subsequent ligand modifications strengthened these interactions, leading to increasingly selective CBP modulators.

**Figure 6.**
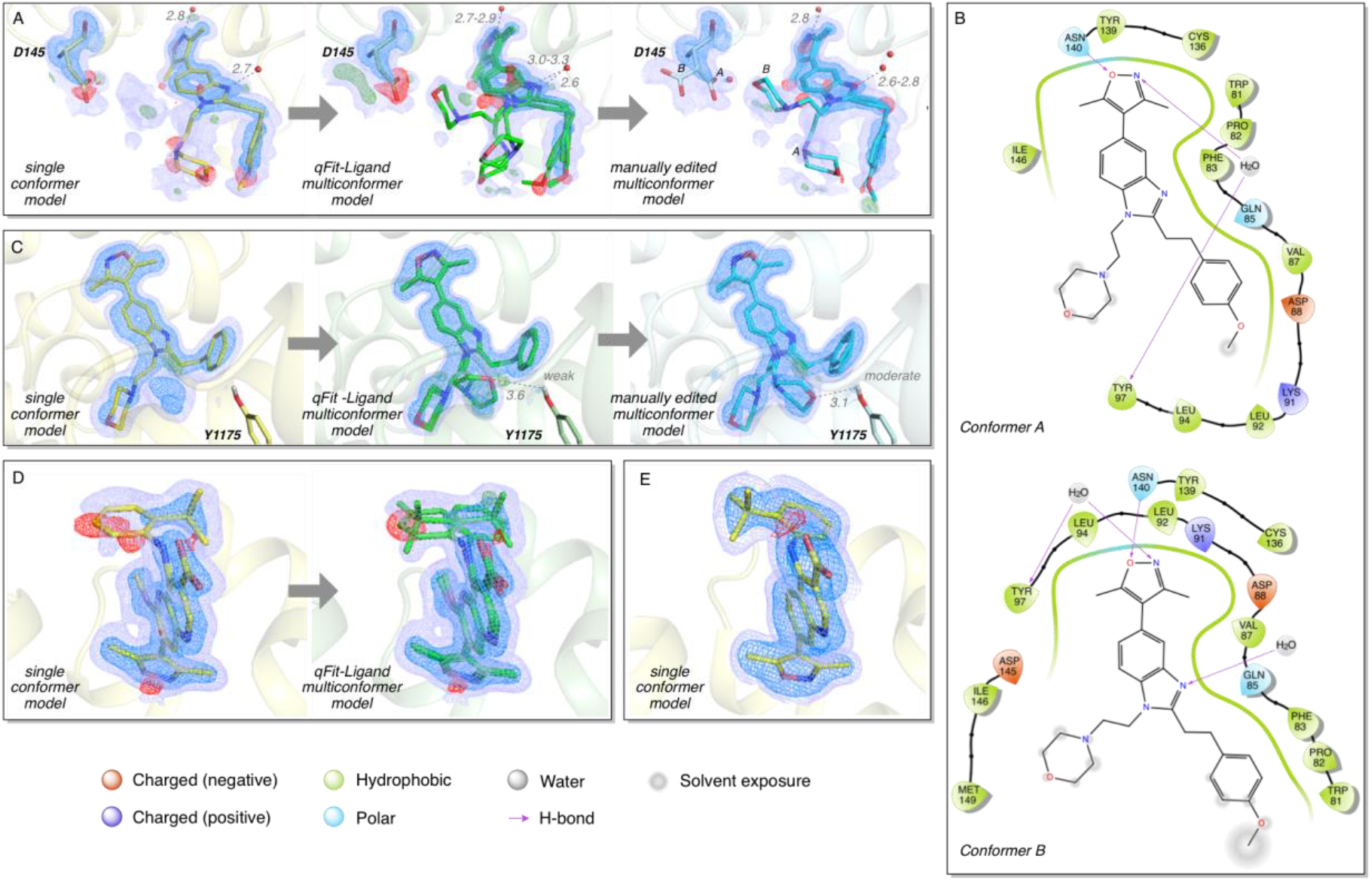
Prospective application of *qFit-ligand* to BRD compounds A) Single conformer crystal structure, *qFit-ligand* multiconformer, and manually edited multiconformer models including alternate conformations of Asp145 and compound BDOIA383 bound to BRD4 (PDB ID 5CFW). B) Protein-ligand interactions of the major, crystal-contact stabilized ‘A’ and minor ‘B’ BDOIA383 conformations of the final *qFit-ligand* model (panel 6A). C) Single conformer crystal structure, *qFit-ligand* multiconformer, and manually edited multiconformer models of a isoxazolyl-benzimidazole ligand bound to CBP BRD (PDB ID 4NR5). D) Single conformer and *qFit-ligand* multiconformer models of ligand 9BM bound to BRD4 (PDB ID 4BW3). Viewing orientation differs from panel A,C to clearly show ligand alternate conformations. E) Ligand S5B’s single binding conformation in BRD2 (PDB ID 4AKN). Electron densities are shown at 1.5σ (blue) and 0.3σ (purple). Positive (green) and negative (red) difference densities are shown at ±3.0σ. Distances in Ångstroms.

Flipped binding modes between BRD4 and CBP had been observed earlier with a bound isoxazolyl-benzimidazole ligand, also leading to improved selectivity.^43^ Strikingly, in this case too *qFit-ligand* identified a minor conformation in the CBP complex structurally close to the BRD4 bound conformation (**Figure 6C**). Because selectivity is often achieved by leveraging unique structural ligand-receptor interactions, identifying alternate ligand conformations may help profile auspicious ligand-receptor secondary molecular recognition sites.

Application of *qFit-ligand* to all 126 BRD2-4 crystal structures in the PDB suggests that differential binding modes and ligand heterogeneity are remarkably ubiquitous (Materials and Methods). Visual inspection and manual curation revealed 12 new binding conformations detected with high confidence (**Figure S7**), and an additional 24 with possible alternate conformations (**Table S5, Figure S6**). The alternate conformation detected by *qFit-ligand* in the crystal structure of BRD4 with ligand 9BM^44^ (phenyl ring flip) is the same conformation observed in the crystal structure of BRD2 with ligand S5B^45^, further supporting the idea that ligands can sample a second, minor pose which becomes dominant after small chemical modifications to their structure. These observations and the pervasiveness of ligand heterogeneity in BRDs suggest that the potential of alternate ligand conformations for structure-based drug design is significantly undervalued.

## Discussion

Ligand conformational heterogeneity is widespread in X-ray data deposited in the PDB, but underreported in the absence of automated and reliable computational methods. q*Fit-ligand* is a new method to model a parsimonious multiconformer ligand in crystallographic electron densities. We formulated the challenge of identifying an ensemble that collectively best agrees with the electron density, from up to tens of thousands of candidate conformations, as a combinatorial optimization problem. *qFit-ligand* relies on exhaustive sampling, iteratively restrained by the electron density, to fully cover the conformational space of each ligand.

We highlighted three examples of previously unmodeled alternate ligand conformations obtained from prospective application of *qFit-ligand* to the D3R and Twilight data sets. Strikingly, even the highly curated D3R data set revealed a previously unmodeled alternate conformation. Targeted application to a single bromodomain receptor suggests that as many as 29 % of receptor-ligand crystal structures could have alternate ligand conformations. This is a likely conservative estimate of the accessible ligand conformational landscape in protein crystal structures. The portrait of rigid receptors and ligands is exacerbated by the common practice of collecting X-ray crystallographic data at cryogenic temperature (100 K). Although cryocooling increases the precision of structure determination by reducing thermal motions, cooling affects the conformational distribution of more than 35 % of side chains in proteins^46^ and has been found to alter ligand binding and abolish transient binding sites observed at room temperature.^47^ Applying *qFit-ligand* to room temperature X-ray crystallography data, which can shift the equilibrium of receptor-ligand conformational ensembles, may reveal additional minor ligand binding poses that are typically masked in cryogenic data.

*qFit-ligand* conformational strain energies after refinement were nearly indistinguishable from those of the manually curated benchmark set, signifying its multiconformer models are chemically accurate. Our analysis showed that *qFit-ligand* conformations were nearly isoenergetic, indicating they could bind in either conformation. Less often we identified a ligand for which the *qFit-ligand* multiconformer model significantly reduced strain energy, suggesting that conformational averaging of the single conformer model had led to a poorly modeled ligand to fit the density. Ironically, multiconformer ligands are commonly filtered out of major test sets for development of docking and conformational sampling approaches, but may be a more ‘trustworthy’ representation of the underlying data^48^. Multiconformer ligand models can therefore address important challenges in ligand validation and deposition in the Protein Data Bank (PDB).^49,50^

Several aspects of *qFit-ligand* could be improved in the future. First, while in principle our conformational sampling approach could be combined with a sophisticated force field,^51^ relying on energy restraints in the discovery stage can result in increased computational cost and can risk excluding promising candidate conformations owing to imperfect sampling and steep potential energy gradients. Rather, we advocate including energy restraints in refinement.^52^ Other conformational search methods with force fields, such as Schrodinger’s ConfGen^53^ and MacroModel,^54^ OpenEye’s Omega,^55^ MOE,^56^ and many other freely available tools^57^ could independently validate results in the absence of data.

Second, the false-positive rates can likely be decreased. Terminal ends and ring flips had low false positive rates, whereas those for the branched and displaced categories were elevated. *qFit-ligand* recovered terminal end and ring flips to within an nRMSD of 0.27. Disordered parts of branched ligands were often solvent-exposed, and therefore more difficult to recover. Displaced ligands are not ‘anchored’ in the binding pocket, and are among the most challenging to recover. An nRMSD of 0.5 or below often sufficed for further refinement, but even nRMSDs up to 0.75 sometimes required only minor manual adjustments, depending on the absolute RMSD between states. For example, ring flips involving rotationally symmetric atom species cannot be uniquely assigned based on the electron density alone, leading to inflated RMSD measures, which are easily adjusted (Materials & Methods, PDB ID 3P4V). All benchmark *qFit-ligand* results were obtained with the same command-line parameters. In practice, specific problems will dictate tailored settings. For example, selecting finer sampling steps or larger volumes for rigid body searches could give better results at the expense of increased computational time. Nonetheless, on our benchmark set, the fully automated *qFit-ligand* multiconformer models were supported by real-space validation measures, and their R_free_ values were statistically indistinguishable from the manually curated set. While these measures cannot perfectly distinguish false positives, structural models consistent with the reflection data create a pool of testable hypothesis, which can be evaluated in drug discovery using ligand structure activity relationships or protein residue mutations.

Third, the partial occupancy of an “unbound”/apo state could be explicitly considered. Weak, overlapping densities originating from partially occupied receptor and ligand conformations in the binding pocket are often difficult to tease apart, owing to a vast number of possible ligand conformations to be evaluated, even in sterically constrained binding cavities. Partially occupied water molecules and crystallographic additives often further confound modeling efforts. In these challenging cases, difference densities from alternate states are often incorrectly resolved by waters. Promising new approaches such as PanDDa^58^ can reveal the electron density of partially occupied states; however, it requires a large number of ‘ground state’ crystal structures to reliably compute their contribution to the partially occupied state. Our method is highly complementary and PanDDa maps could even be used as input to *qFit-ligand*. Synergy of these approaches holds the promise to enable efficient, accurate and unbiased discovery of alternate fragment and ligand binding poses. This is increasingly important in view of the ability to structurally screen hundreds of candidate ligands within hours on modern synchrotrons. The combination of these methods may help remove temptation to fill all difference density with waters, while avoiding the overly optimistic modeling of partial occupancy ligands that are highlighted by the Twilight database.

Finally, as the particle size and resolution limitations of single-particle cryo-electron microscopy (cryo-EM) continue to improve, that technology will have a major impact on drug discovery.^59^ High-throughput and automation approaches,^60^ combined with the size of the complexes in cryo-EM structure determination, will soon turn careful modeling of protein and ligand structural heterogeneity into a major bottleneck. *qFit-ligand* can provide an efficient, automated modeling approach at the amino-acid length-scale, as EM maps are immutable during modeling and refinement. Beyond applications to drug discovery, as time-resolved serial crystallography is rapidly becoming routine at X-ray free electron lasers and even synchrotrons,^61^ *qFit-ligand* can help resolve minor populations of structural protein-ligand intermediates in light-driven pump-probe^62^ or structural enzymology ‘mix-and-inject’ experiments.^63,64^

Revealing the full receptor-ligand conformational ensemble can help drug design by exploiting the balance between entropy and enthalpy in compound design^65,66^ and by characterizing the effect of pre-rigidifying ligands on affinity.^67^ Equally important, it can help rational design of ligand selectivity by exposing accessible molecular surfaces unique to their intended targets.^68^ In the future, full integration of *qFit-ligand* with *qFit* could reveal the structural reorganization of binding pockets and allosteric signal propagation in the receptor upon ligand binding.^18,69^ Our *qFit-ligand* open source software, available from https://github.com/ExcitedStates/qfit_ligand, provides promising, new starting points for ligand optimization and structure based drug discovery. However, communication between structural biologists, computational chemists, and medicinal chemists remains a requisite for successful, rational design.

## Materials and Methods

### Survey of the Protein Data Bank and benchmark creation

All structure coordinate files were downloaded from the PDB. Structures determined by X-ray crystallography were checked for *HETATM* entries (ligands) containing at least two different *altloc* identifiers with identical chemical composition. Ligands with fewer than 15 non-hydrogen atoms and covalently linked ligands were discarded. This resulted in a list of 2,611 ligands divided over 1,845 PDB files. The list was further pruned to exclude ligand flips, i.e. alternate conformations that do not have a common cluster of atoms in space, and alternate conformations consisting exclusively of ring puckers as our algorithm was not designed to sample these types of conformational changes. We then selected receptors and ligands of pharmaceutical interest, resulting in a final benchmark set of 90 cases that refined against the deposited structure factors and CIF files. Each case was re-refined using phenix.refine v1.11 with the following parameters

#### For PDBs better than 1.5Å

optimize_xyz_weight=true optimize_adp_weight=true optimize_mask=True main.number_of_macro_cycles=10 adp.individual.anisotropic=“not water and not element H” adp.individual.isotropic=“water or element H”

#### For PDBs worse than 1.5Å

optimize_xyz_weight=true optimize_adp_weight=true optimize_mask=True main.number_of_macro_cycles=10 adp.individual.isotropic=all

Single conformer ligand models were created by removing the ligand’s ‘B’ conformation from the original deposited PDB model and resetting the occupancy of the ‘A’ conformation to 1. The single conformer models were re-refined as above.

### The *qFit-ligand* approach

The *qFit-ligand* algorithm takes as input the initial structure of a single conformer ligand modeled in the electron density, a real-space map (2mF_o_-DF_c_) in *ccp4* format and its resolution, and, optionally, the receptor and other ligand and solvent atoms for clash avoidance. The *qFit-ligand* algorithm starts by scaling the map electron density values to approximate absolute scale, using only the density under the footprint of the receptor. Next, the algorithm calculates an F_c_ map corresponding to the receptor and any other atoms in the crystal not part of the ligand. This map is subtracted from the experimental density. Map values below the mean are set to zero to prevent building into spurious density.

Next, *qFit-ligand* determines rotatable covalent bonds of the ligand and rigid groups of atoms. Rigid groups of atoms do not have internal, rotatable covalent bonds, such as terminal atoms or ring systems. Covalent bonds are determined based on proximity; i.e., if the distance between two atoms is smaller than their combined covalent radius plus 0.5Å, the atoms are considered covalently bonded. A covalent bond is rotatable if it is not part of the same ring system. Hybridization states are ignored in the current implementation.

After preparing the input density and ligand, *qFit-ligand* alternately exhaustively samples the rotatable bonds of the ligand and determines the optimal occupancy of each conformation. To prevent a combinatorial explosion, the ligand is iteratively built up starting from each rigid group of atoms. The first sampling iteration consists of a local rigid body search of the starting group within a box with an edge size of 0.4 Å at a 0.1 Å interval and 10 randomly generated orientations at a maximum rotation angle of 10 degrees (1,250 conformations). The following iterations each sample N (default 2) torsion angles using a pre-set (default 6°) step size. Clashing conformations, both internal and with the receptor (using an efficient, **O**(1), spatial hashing algorithm), are detected and removed from the set. Bond lengths and angles are kept fixed during the whole procedure.

After each sampling iteration, the optimal occupancies of generated conformations are determined as follows. Each conformation is transformed into a density given by

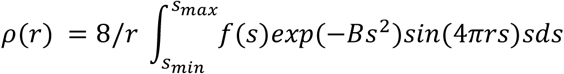

where *r* is the position compared to the atom position, *f(s)* is the atomic scattering factor as a function of the momentum vector of the atom, *B* is the isotropic B-factor, and *s_min_* and *s_max_* are the minimum and maximum momentum vector length. For computational efficiency a lookup table is created for each atom at an interval of 0.01 Å.

A combined mask is calculated by forming the union of all individual conformation masks, using a resolution dependent radius, where *r* = 0.5 + R / 3. Density values under the footprint of the resulting mask are extracted and used as input for Quadratic Programming (QP) and Mixed Integer Quadratic Programming (MIQP) to select up to 5 conformations that best represent the density data locally. QP and MIQP solvers guarantee the global optimal occupancy of each conformation by minimizing the real-space residual given by

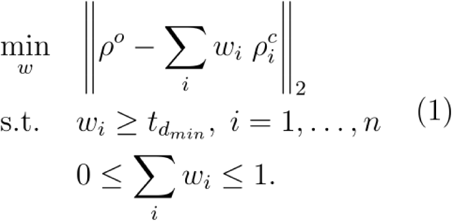

where *ρ*^o^is the 2mF_о_-DF_c_ map, *ρ*_*i*_ is the calculated density of conformer *i*, *w_i_* is the weight, or occupancy, of conformer *i*, and 0 ≤ *t* ≤ 1 is a threshold on the occupancy. If *t* = 0, the constraints enforce non-negativity for the occupancies, and the program is a QP. The combined occupancy cannot exceed unity. Selecting t > 0 introduces sparsity or threshold constraints, which turns the program (1) into a MIQP. To reduce computational complexity, relevant conformations are preselected using QP and used as input for a subsequent round of MIQP.^20^

After the ligand has been fully built up starting from each cluster, all resulting conformations are pooled together for a final round of rescoring. The individual Pearson product-moment cross-correlation (RSCC) is calculated for each individual conformation. Conformations with RSCC score less than 0.9 times the highest correlation, and redundant conformations for which all pairwise atom distances are within those of previously accepted conformations, are discarded. Finally, a parsimonious multiconformer model is created as follows. An MIQP step at an occupancy threshold of 0.20 is performed on the remaining pool of conformations, the resulting selected conformations are ordered by RSCC value, and starting from the conformation carrying the highest RSCC, additional conformations are added if the RSCC increases under the combined footprint and else discarded. This is performed iteratively until self-consistent, i.e. all conformations increase the RSCC under the combined footprint. The output of *qFit-ligand* consists of all conformations before the final rescoring round and the sparse occupancy-weighted multiconformer ligand model.

*qFit-ligand* is implemented in Python 2.7 and relies on the open-source NumPy, SciPy and CVXOPT^70^ packages and the freely available Community Edition of IBM ILOG CPLEX, with added modules from the mmLib Python toolkit.^71^ *qFit-ligand* is released under the MIT license and can be downloaded free of charge from https://github.com/ExcitedStates/qfit_ligand, where additional documentation and installation instructions can be found.

### Benchmarking *qFit-ligand*

Simulated structure factors were generated from the re-refined benchmark set using *phenix.fmodel* at resolutions of 1.0, 1.3, 1.6, 2.0 and 2.5 Å resolution. Occupancy of the B-conformer was varied from 0.5 to 0.0 occupancy in 0.1 decrements, requiring that the combined occupancy of the multiconformer ligand summed up to unity. A random error of 10 % was added to the amplitudes, a fraction of 0.1 was used for R_free_ flags, and we selected k_sol_ = 0.4 and b_sol_ = 45Å^2^. The B-factor of each ligand atom was adjusted by 10 times the difference in real and simulated resolution, i.e. the B-factor was inflated for lower resolution simulated data and sharpened for high resolution data. The resulting *mtz* files were converted into *ccp4* density files using *phenix.mtz2map.*

For each resolution and occupancy combination, *qFit-ligand* was run on the whole benchmark using different sampling parameters: sampling 1 degree of freedom (DoF) using a torsion sampling interval of 1 degree, and sampling 2 DoFs simultaneously using a torsion sampling interval of 6 degrees. The results were analyzed by calculating a normalized RMSD

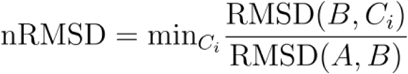

between the output structures *C_i_* of *qFit-ligand* and the ligand *B* conformer used during the structure factor generation, normalized by the RMSD between the A and B conformer. Conformers *C_i_* for which nRMSD < 0.5 are more similar to the B conformer than the A conformer. We note that the RMSD (and therefore also the nRMSD) measure has important limitations, affecting the results. For example, since RMSD is typically calculated between unique points, it fails to account for symmetric transformations within a ligand, e.g. a 180° flipped acidic group or aromatic ring, thus introducing additional error. Unfortunately, to our knowledge there is no readily available method that addresses this issue. While a low nRMSD guarantees a good solution, a relatively high nRMSD can still suggest an alternate conformation that requires some manual adjustments, including symmetric transformations.

We subsequently evaluated the performance of *qFit-ligand* for each category in the benchmark set, at simulated resolutions and occupancies, after refinement with PHENIX v1.11. The final selection stage was optimized heuristically by finding a balance between maximizing sensitivity and minimizing false-positives against the benchmark.

### D3R and Twilight database investigation

We downloaded the D3R (https://drugdesigndata.org/about/datasets) and Twilight (http://www.ruppweb.org/twilight/newligands-2016.tsv.bz2) data bases deposited in 2016. For all cases, structure coordinates and 2mF_o_-DF_c_ maps were downloaded from the PDB. For the Twilight cases, we discarded structures with a reported resolution worse than 2.0Å and ligands consisting of less than 15 and more than 35 non-hydrogen atoms and a cross-correlation score less than 0.6. The resulting *qFit-ligand* multiconformer models were ranked on the Fisher ztransformation score^72–74^, which we apply to crystallography data for the first time. Starting with the conformation with the highest RSCC, we calculated

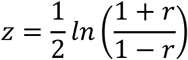

where *r* is the RSCC, and its associated standard error

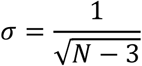

where *N* represents the number of independent observations, approximated by *N* = *MV*/*R*, where *MV* represents the molecular volume of the ligand and *R* the resolution. As additional conformations are added, the Fisher z-transformation score is recalculated for both the starting conformation and combined multiconformer under the footprint of the latter. The resulting difference in z-score is divided by the standard error to provide a resolution and size corrected measure of the increase in cross-correlation, where higher is better. The highest z-score difference found over all added conformations is reported and ranked accordingly.

### Ligand Energy

To access ligand conformational energies, we carried out constrained minimizations (flat bottom width of 0.2Å) of all ligands (from single conformer and *qFit-ligand* multiconformer models) using Jaguar^75^ with the M06-2X functional^76,77^ and 6-31G(d,p) basis set, except for bromine atoms which used the LAV2P basis set. Gas phase energies of ligand conformations generated by *qFit-ligand* (which were subsequently refined) were compared to ligand conformations in the single conformer model (i.e., 0 kcal/mol). The relative energies of each alternative conformation in the *qFit-ligand* multiconformer model were multiplied by their respective occupancies, then summed to arrive at the occupancy-weighted ligand energy.

## Acknowledgements

We gratefully acknowledge support from Relay Therapeutics. D.A.K. was supported by an A.P. Giannini Postdoctoral Fellowship. R.F. is supported by a Novo Nordisk Foundation and Stanford Bio-X Program fellowship NNF15OC0015268. J.S.F. is supported by a Searle Scholar Award from the Kinship Foundation, a Pew Scholar Award from the Pew Charitable Trusts, a Packard Fellowship from the David and Lucile Packard Foundation, NIH GM123159, NIH GM124149, UC Office of the President Laboratory Fees Research Program LFR-17-476732, and NSF STC-1231306. H.v.d.B. is supported by NIH GM123159.

## Author Contributions

GCPvZ, RF, AH, HvdB developed and implemented core algorithms; GCPvZ, BMH, KB, TD, JSF, HvdB analyzed data; DAK, PS contributed code and data analysis; JSF and HvdB conceived the study. GCPvZ, BMH, JSF, and HvdB wrote the manuscript. All authors edited the manuscript.

## Supplementary Figures

**Figure S1.**
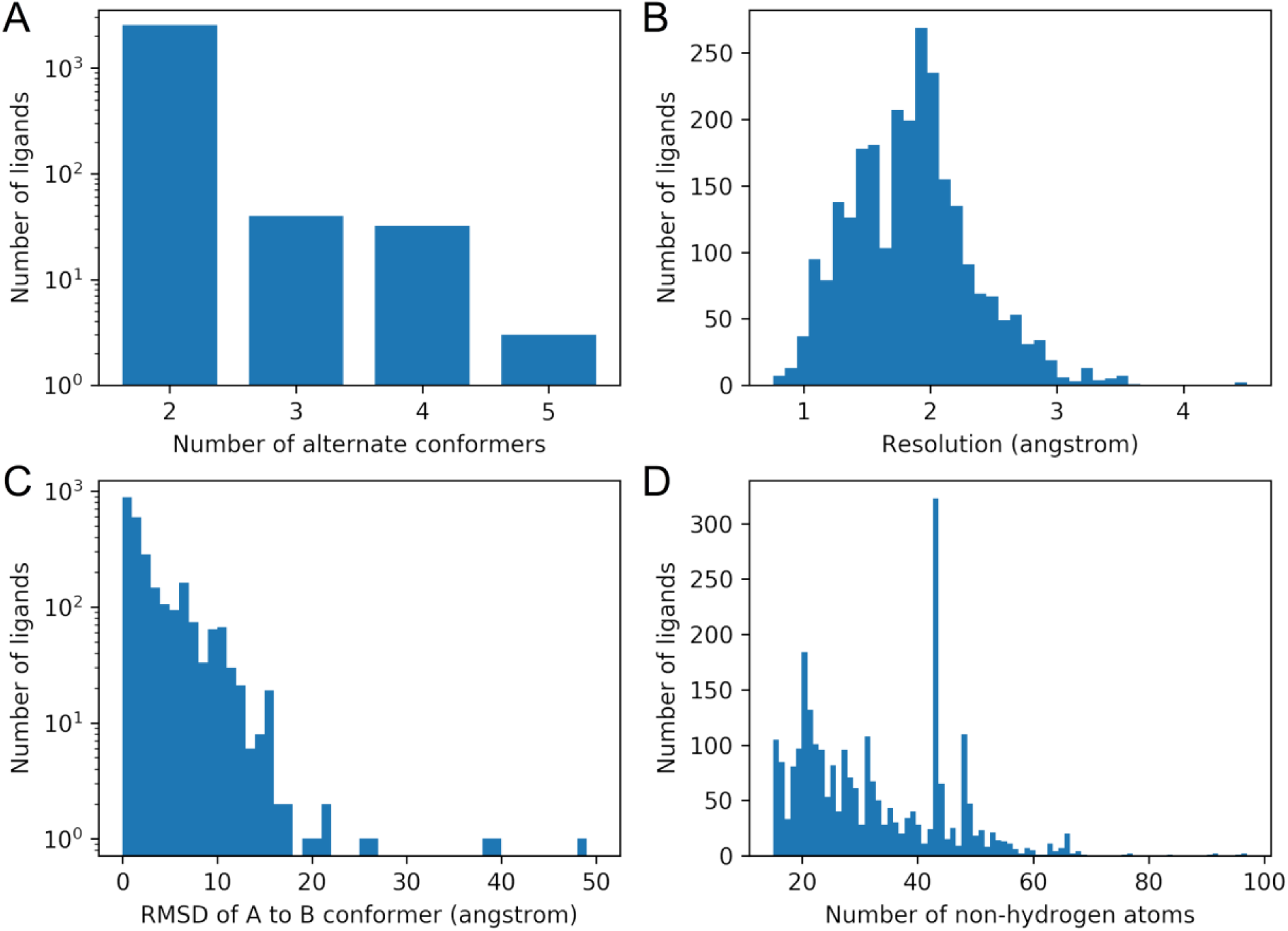
Overview of 2,611 multiconformer ligands deposited in the PDB as of June 2017. The most common ligands found were heme (240), NADP (48), and NAD^+^ (46), while most other ligands only appear once (714) or twice (207 unique codes). A) Histogram of number of alternate conformers per ligand. The majority have two conformers (2,534), while less than 100 have three to five conformers. B) The resolution of the multi-conformer ligands ranges from 0.5Å to 4.5Å, with the mode at 2.0Å. C) Histogram of RMSD between A and B conformer. Large RMSD values indicate alternate binding pockets instead of alternate conformations. D) Distribution of ligand size by the number of non-hydrogen atoms.

**Figure S2.**
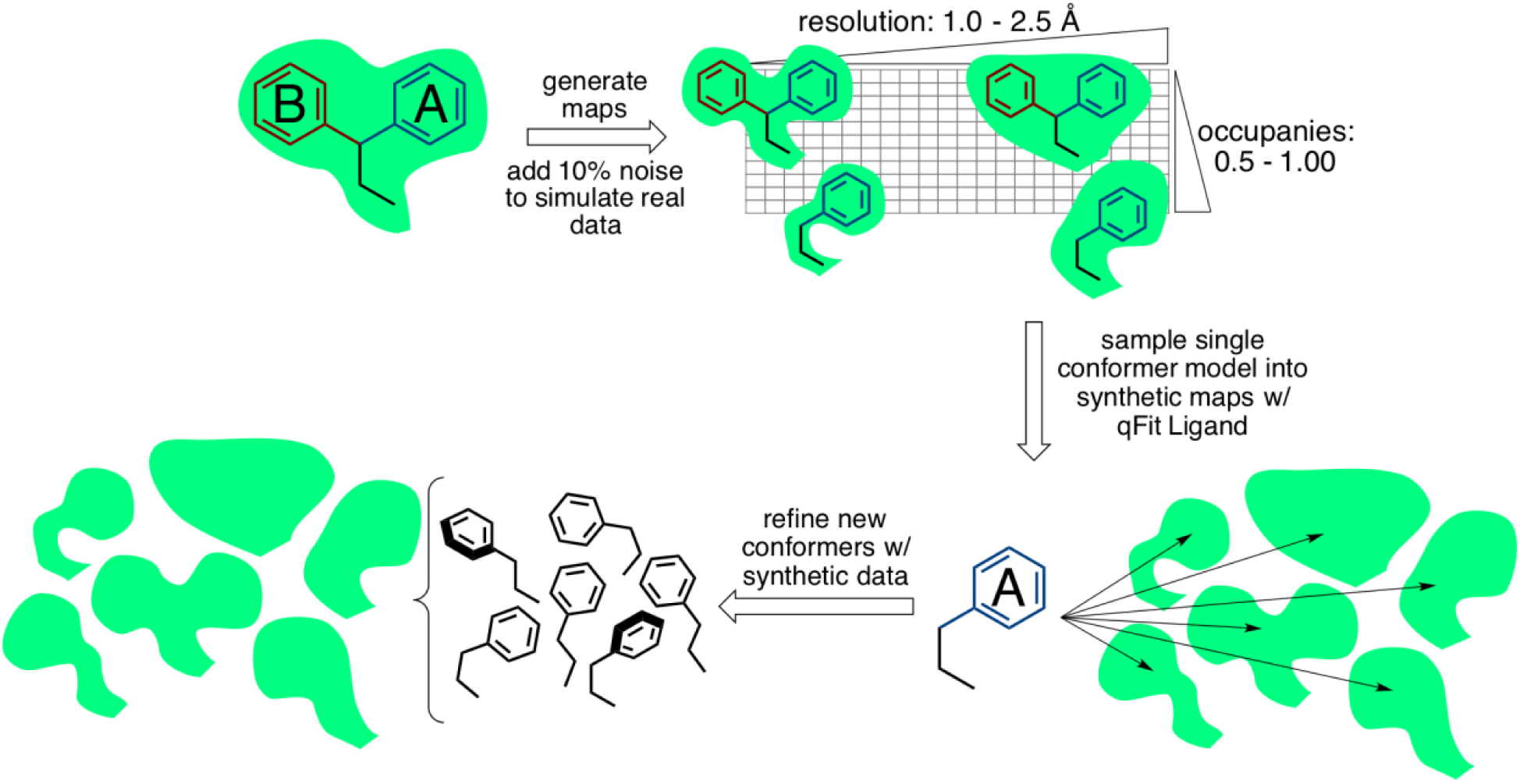
Generating simulated data for testing resolution and occupancy limits of *qFit-ligand*.

**Figure S3.**
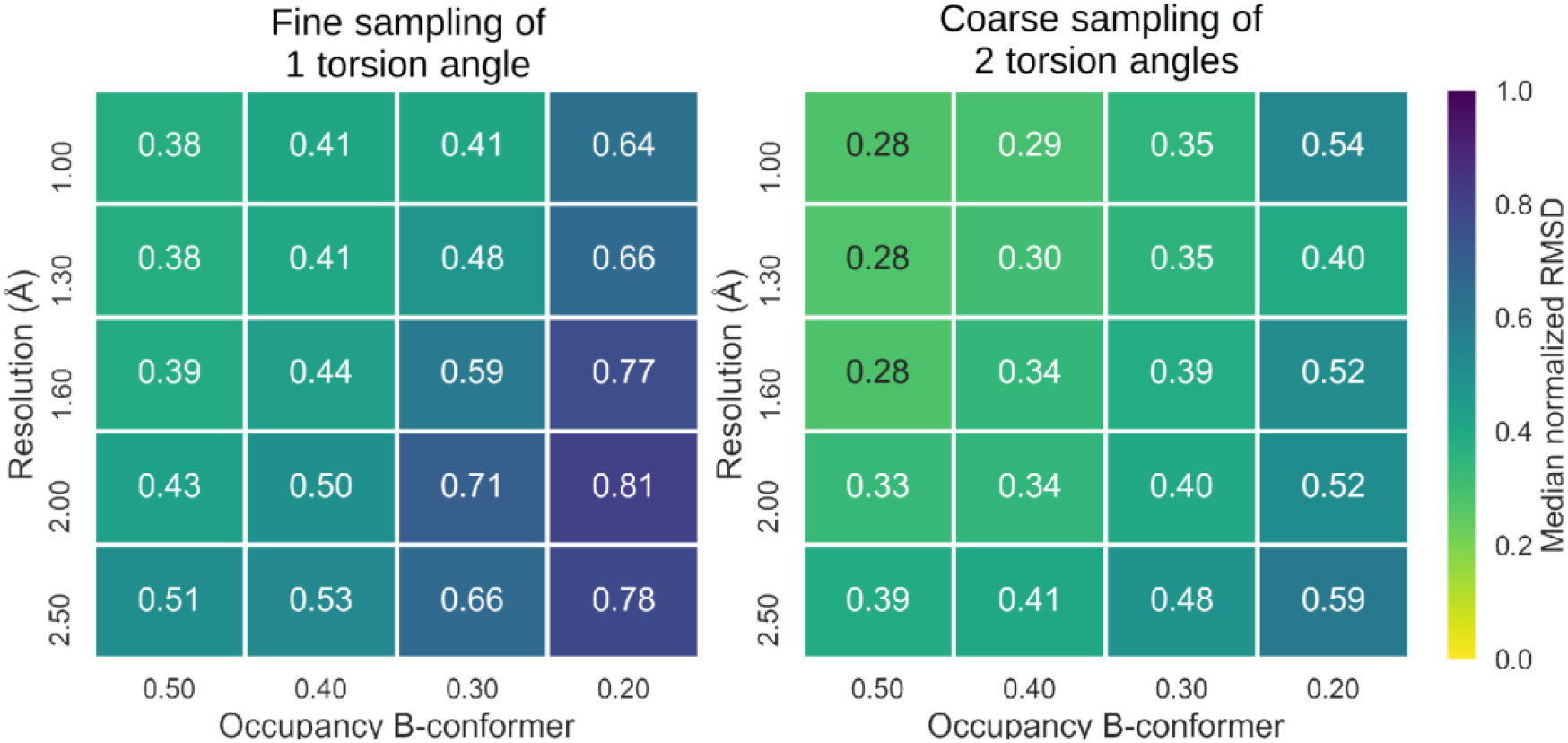
Performance of *qFit-ligand* before final rescoring round measured in median normalized RMSD when exhaustively sampling 1 torsion angle at a fine interval of 1 degree (left) and 2 torsion angles at a coarser interval of 6 degree (right).

**Figure S4.**
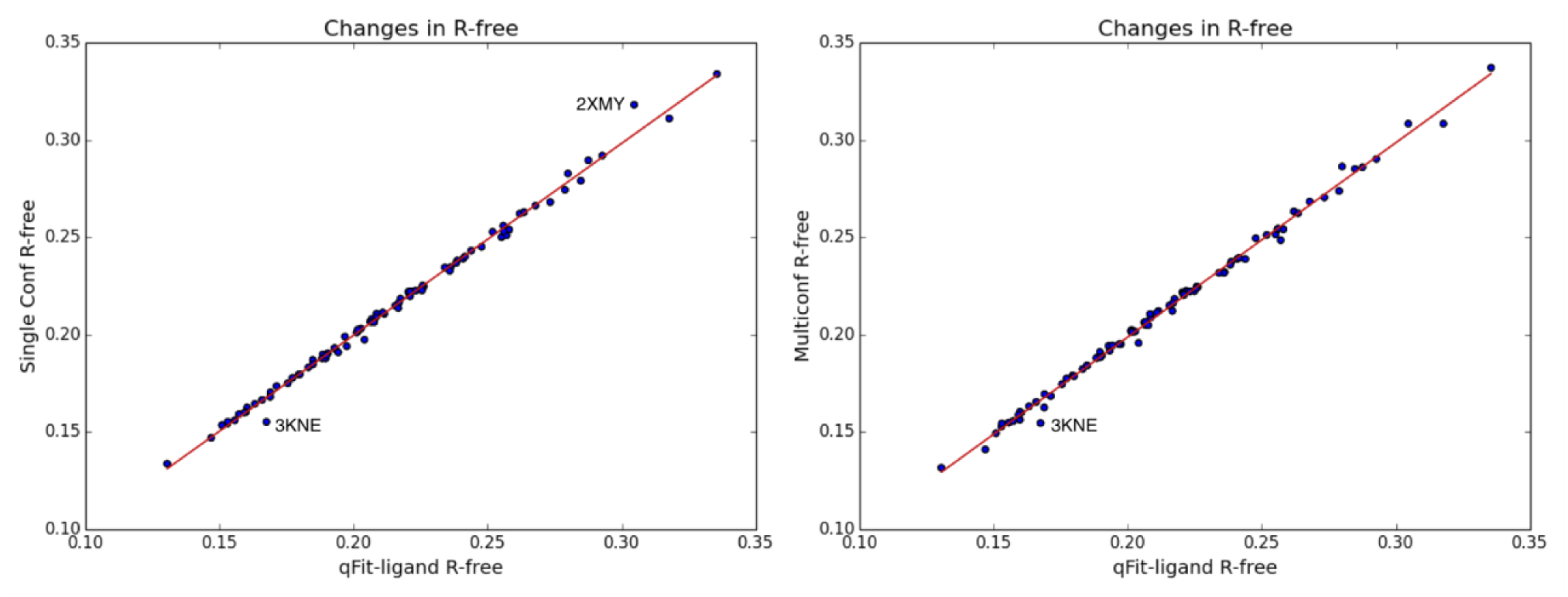
R-free values post-refinement of *qFit-ligand* multiconformer model vs. single conformer and deposited multiconformer models, respectively, from our benchmark dataset. Outliers are labeled.

**Figure S5.**
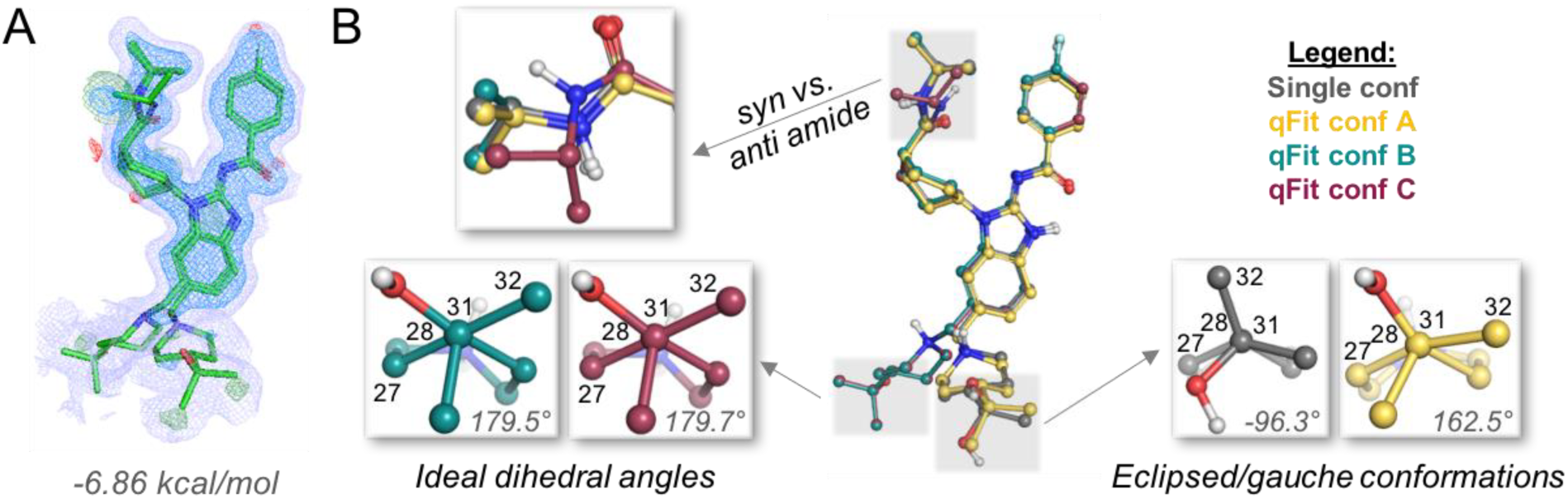
Branching ligand heterogeneity from the benchmark dataset (PDB ID 4FOD). A) Ligand conformations generated by *qFit-ligand* are significantly lower in energy (by ∼7 kcal/mol) than the corresponding single conformer model. Electron densities are shown at 1.5σ (blue) and 0.3σ (purple). Positive (green) and negative (red) difference densities are shown at ±3.0σ. B) Evaluation of ligand conformations reveal that an *anti* amide conformation and ideal dihedral angles as opposed to *syn* amide conformation and eclipsed/gauche conformations help to significantly lower ligand conformational energies.

**Figure S6.**
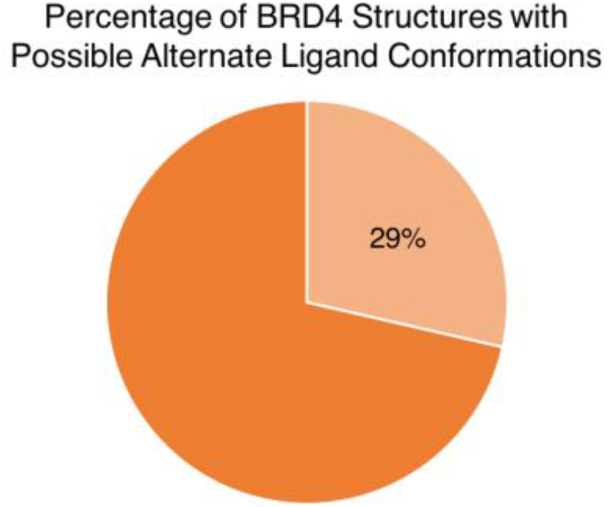
Possible alternate ligand conformations found in BRD4 structures. *qFit-ligand* was applied to a dataset of 126 crystal structures (100 % sequence similarity to 5CFW) deposited in the PDB. Of those, *qFit-ligand* detected alternate conformations in 12 crystal structures with high confidence, and another 24 with possible alternate ligand conformations.

**Figure S7.**
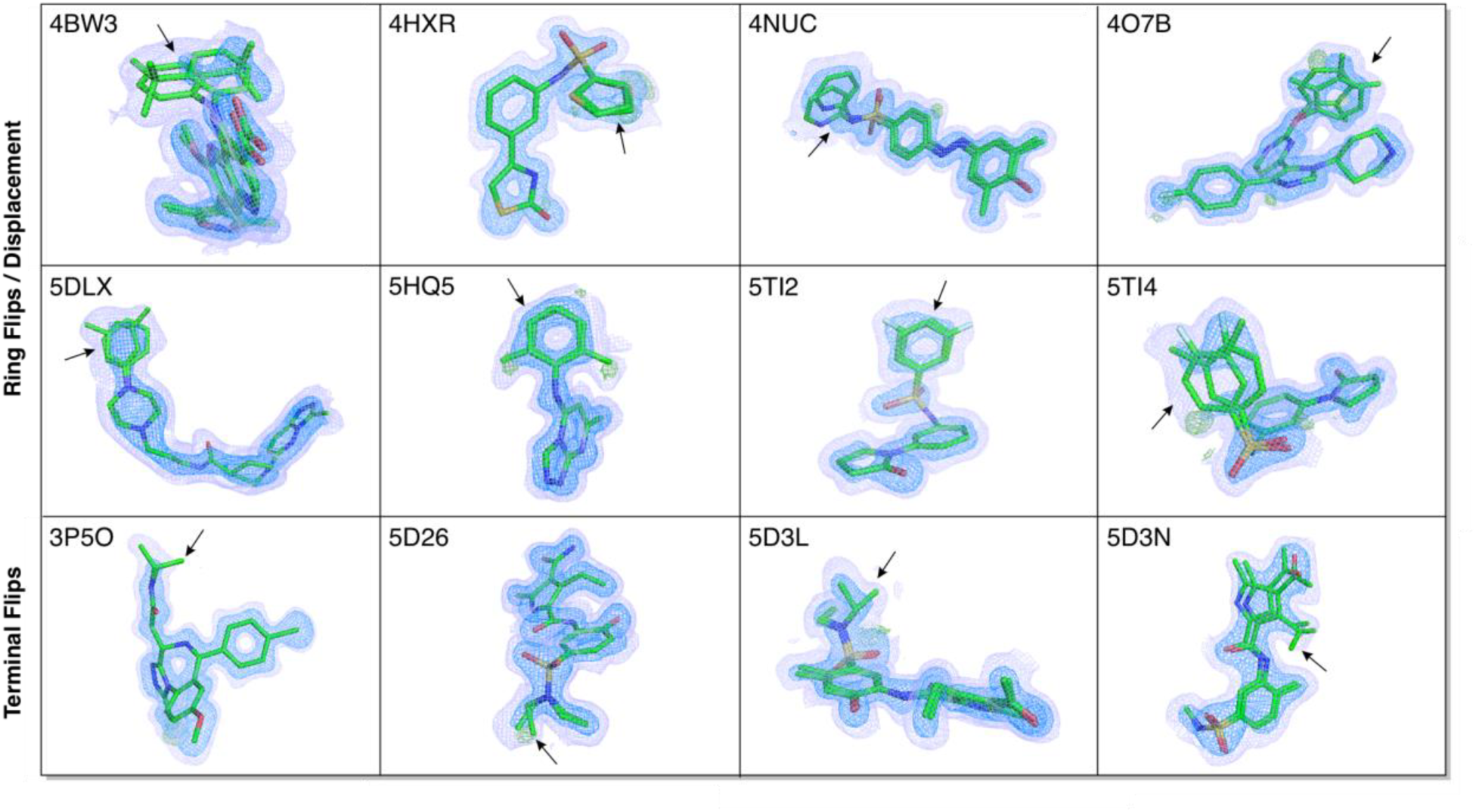
Twelve new binding conformations detected with high confidence for ligands bound to BRD4. Electron densities are shown at 1.5σ (blue) and 0.3σ (purple). Positive (green) difference densities are shown at ±3.0σ.

**Figure S8.**
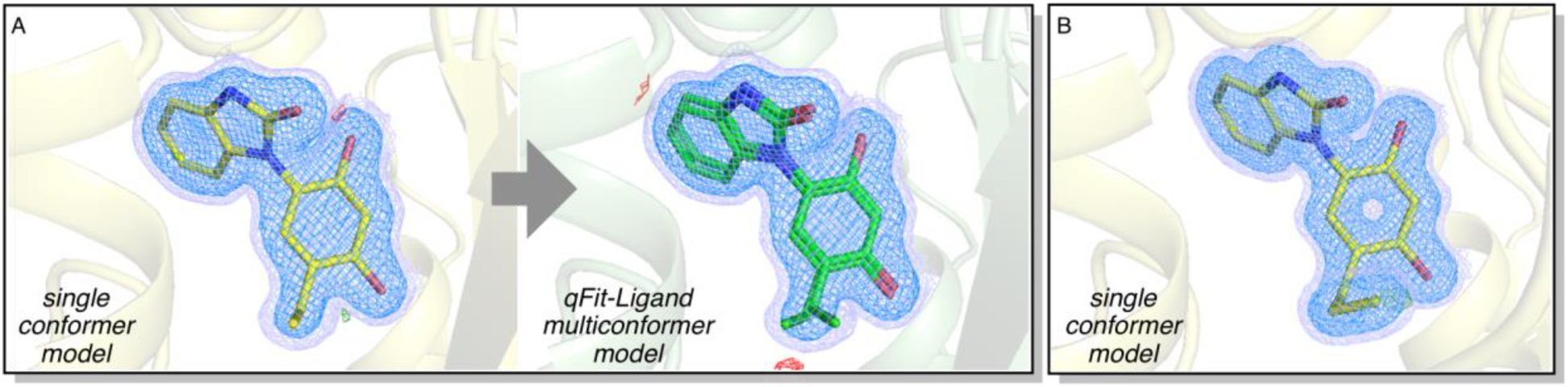
Prospective application of *qFit-ligand* to inhibitor CS301 bound to HSP90 (PDB ID 4YKQ) from the D3R dataset. A) Two discrete conformations of the ethyl group is found by *qFit-ligand*. This terminal flip could be used as a design element for future ligands; for example, we imagine a derivative of the ligand with a cyclopropyl group in place of the ethyl group. B) Although this derivative was not synthesized, a derivative of inhibitor CS301 with a propyl group was found to occupy the space captured by our qFit-ligand multiconformer model, providing further evidence of the alternate conformation shown in panel A. Electron densities are shown at 1.5σ (blue) and 0.3σ (purple). Positive (green) and negative (red) difference densities are shown at ±3.0σ.

**Figure S9.**
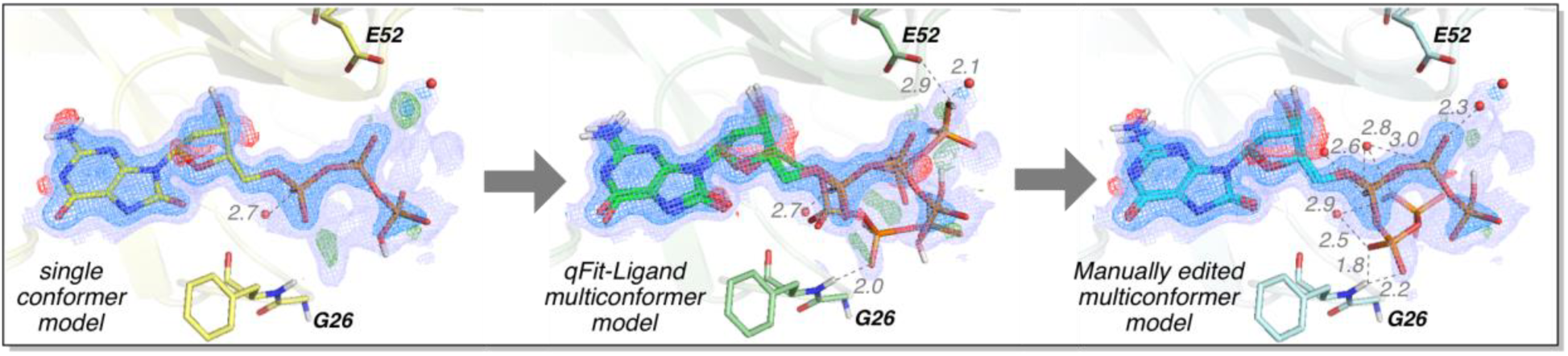
Prospective application of *qFit-ligand* to substrate 8-oxo-dGTP bound to MTH1 (PDB ID 5FSI). *qFit-ligand* multiconformer, and manually edited multiconformer models shown. In the crystal structure, the electron density provides evidence of ligand disorder. Two of the alternate conformations of the phosphate tail chosen by qFit-ligand better explain the ligand density and, furthermore, reveals protein-ligand interactions that were not captured in the single conformer model Single conformer, Electron densities are shown at 1.5σ (blue) and 0.3σ (purple). Positive (green) and negative (red) difference densities are shown at ±3.0σ. Distances in Ångstroms.

**Figure S10.**
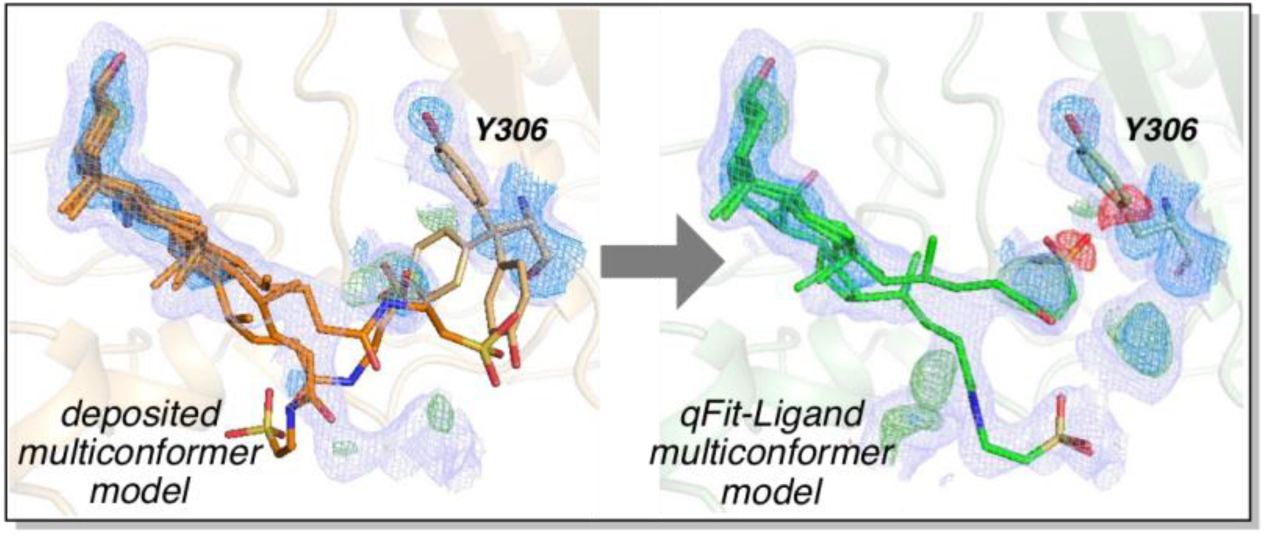
Prospective application of *qFit-ligand* to an ENPP2-bound ligand (PDB ID 5DLV), which includes multiple conformations of the ligand and Y306. Upon inspection of the density, we removed two of the three Y306 and ligand conformations. Deposited multiconformer model and *qFit-ligand* multiconformer model shown. *qFit-ligand* conformations fit the density better than the manually curated, deposited multiconformer model. The presence of positive difference densities are likely a result of unmodeled waters. Electron densities are shown at 1.5σ (blue) and 0.3σ (purple). Positive (green) and negative (red) difference densities are shown at ±3.0σ. Distances in Ångstroms.

**Table S1.** Overview of benchmark cases and categorization of conformational change between alternate conformers.

**Table S2.** Performance of *qFit-ligand* on the benchmark using re-refined deposited data, showing quality metrics of re-refined single conformer structures, re-refined multiconformer deposited structures and refined *qfit-ligand* generated structures. The single conformer structures were generated by removing the ligand’s B-conformer from the deposited data with subsequent refinement.

**Table S3.** Results of applying *qFit-ligand* on a subset of the Twilight database. Entries are sorted by the highest found Fisher z-score. The *Twilight score*, R_work_ and R_free_ were retrieved from the Twilight database. Entries in the *RMSD A ‐ > B (Å) (deposited)* column are non-empty if the ligand in the deposited PDB file has an alternate conformer modeled.

**Table S4.** Overview of prospective cases with an unmodeled alternate conformer found by *qFit-ligand*.

**Table S5.** BRD4 structures subjected to *qFit-ligand* approach, ordered by the Fisher z-score.

